# Rapid and dynamic nucleic acid hybridization enables enzymatic oligonucleotide synthesis by cyclic reversible termination: A novel mechanism for enzymatic DNA synthesis

**DOI:** 10.1101/561092

**Authors:** K. Hoff, M. Halpain, G. Garbagnati, J. Edwards, W. Zhou

## Abstract

Phosphoramidite chemistry for DNA synthesis remains the industry standard despite limitations on length and yield of the resulting oligonucleotides, time restrictions, and the production of hazardous waste. Herein, we demonstrate the synthesis of single-stranded oligos on a solid surface by DNA polymerases and reverse transcriptases. We report single base extension of the surface-bound oligonucleotide which transiently hybridizes to a neighboring strand with as few as the last two bases. Additionally, when multiple transient hybridization structures are possible, each templating a different base, a DNA polymerase or reverse transcriptase can extend the oligonucleotide with either of these two bases, and therefore the sequence of the newly synthesized fragment can be controlled by adding only the desired base (dNTP deoxyribonucleic acid triphosphate) to create custom oligonucleotides. We used this enzymatic approach to synthesize a 20 base oligonucleotide by incorporating reversible terminator dNTPs through a two-step cyclic reversible termination process with stepwise efficiency over 98%. In our approach, a nascent DNA strand that serves as both primer and template is extended through polymerase-controlled sequential addition of 3’-reversibly blocked nucleotides followed by subsequent cleavage of the 3’-capping group. This process enables oligonucleotide synthesis in an environment not permitted by traditional phosphoramidite methods, eliminates the need for hazardous chemicals, has the potential to provide faster and higher yield results, and synthesizes DNA on a solid support with a free 3’ end.

## Introduction

The first chemically synthesized dinucleotide, dTdT, was reported in 1955(1), and the first gene was synthesized in 1970(2). This gene, encoding a transfer RNA, was just 77 bases long and synthesized by enzymatically joining 17 chemically synthesized short oligonucleotides (2). While today, anyone can purchase custom oligonucleotide sequences up to several thousand bases long, the demand for longer and more accurate synthetic DNA is increasing as DNA and RNA are used for new applications in therapeutics (3,4), high-throughput genotyping (5), gene and whole genome synthesis(6), and data storage (7). These new applications place high requirements on stepwise yields, but chemical synthesis methods can only achieve 98.5-99.5% stepwise efficiency. For example, the final yield for a 1kb fragment synthesized with 99.5% efficiency per step is less than 1%. Therefore, long sequences are enzymatically assembled from shorter, purified, chemically synthesized fragments, and they are then enzymatically processed and screened to remove inaccurate sequences or to correct for errors. This process can be arduous and the ability to synthesize longer, more accurate sequences would enable emerging advanced technologies limited by this multi-step method(6).

The first published Enzymatic Oligonucleotide Synthesis (EOS) methods used RNases (8) and polynucleotidyl phosphorylase(9). Oligonucleotides synthesized with these enzymes enabled research that was otherwise impossible at the time(10), but the methods were highly inefficient compared with emerging phosphoramidite chemistry and subsequently abandoned. Attempts at EOS have also been made with ligases(11), but all recent published efforts have utilized Terminal deoxynucleotidyl Transferase (TdT). TdT is a unique DNA polymerase that can extend single-stranded DNA but not a DNA duplex due to a 16 amino acid lariat-like loop that occupies the binding site for a templating DNA strand(12). Several enzymatic oligonucleotide synthesis approaches using TdT have been attempted; cyclic reversible termination with 3’-blocked nucleotides(13–15), using a reversible covalent TdT-nucleotide complex (16), or simply using dNTPs to synthesize homopolymer tracts for data encryption(17,18). For various reasons, these methods have experienced limited success, and the longest product accurately synthesized using TdT is just 10 bases. However, as the demand for longer, more accurate, less expensive oligonucleotides increases, polymerase-mediated methods are a natural direction to explore, because *in vivo*, these enzymes are capable of high-fidelity synthesis of an entire genome(19). Therefore, we sought to develop a cyclic reversible termination method using a polymerase that efficiently incorporates 3’-blocked reversible terminators, and herein we demonstrate controlled synthesis of single-stranded 20mer DNA oligonucleotide with stepwise efficiency over 98%. Ultimately, we envision a path forward using our methods to achieve efficiencies that exceed traditional phosphoramidite methods.

## Results

### Transient Secondary Structures in DNA Enable Synthesis

Herein, we demonstrate the unique ability of some DNA polymerases and reverse transcriptases (RTs) to extend single-stranded DNA, as shown in Fig. 1a. Since the enzymes tested lack terminal deoxynucleotidyl transferase activity, we hypothesized that the polymerase extends the oligonucleotide when it forms a transient hybridized structure though the sequence of the oligonucleotide used can only hybridize two bases at the 3’ end. We sought to harness this capability for EOS, and we generated a series of modified 9°N DNA polymerases capable of efficiently incorporating reversible terminators in these reactions, which we refer to as Duplases. The oligonucleotide extension products are templated single base extensions that are the result of transient hybridization, and Duplase extension proceeds in a sequence-specific manner with only two bases of transient hybridization which may be occur internally or opposite a neighboring strand (Fig. 1b-c). Although extension with a nucleotide not templated by a neighboring sequence can be observed at longer timepoints, we hypothesize that this is the result of misincorporation, and for shorter one-minute reactions, complete incorporation with either of two the templated base is observed, and no incorporation is observed with the other two non-templated bases (Fig. 1b).

**Fig. 1.**
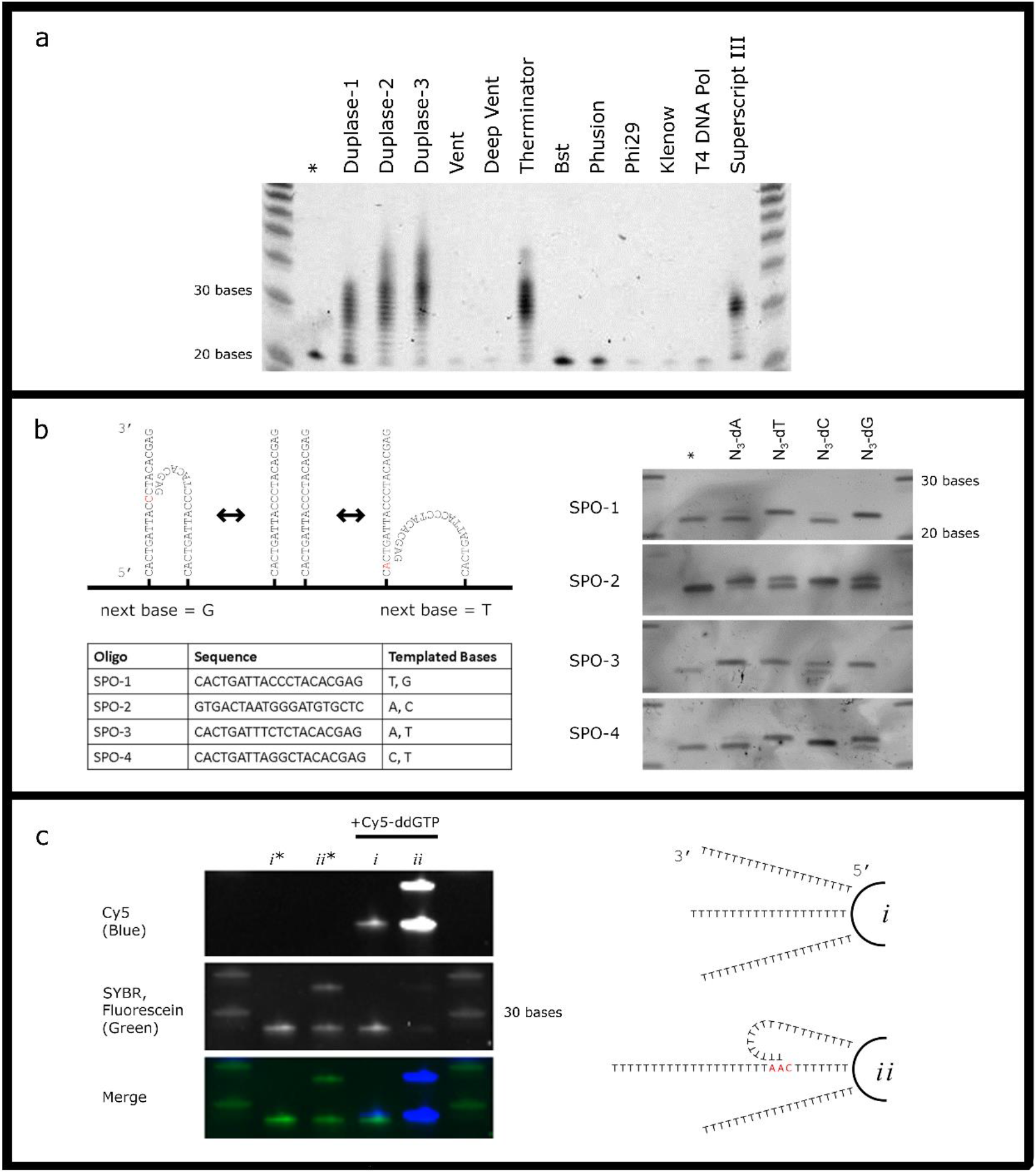
Dynamic hybridization of DNA enables EOS. (a) **Extension of single-stranded DNA by DNA polymerases and reverse transcriptases.** Denaturing PAGE analysis of a 20 base single-stranded sequence, self-priming oligo-1 aka SPO-1, using different enzymes and dGTP. Despite a maximum of two bases of hybridization, some enzymes are able to extend this solid-phase oligonucleotide. None of these enzymes have previously been reported to have nucleotidyl transferase activity on single-stranded DNA. **(b) Sequence-specific extension of single-stranded DNA.** Duplase extension of four different 20 base oligos proceeds in a sequence-specific manner, and only two bases of hybridization are required for extension. * indicates unextended control. Extension with non-templated bases can be explained by misincorporation that occurs at long reaction times**. (c) Extension of solid-phase oligos through intermolecular reactions.** Magnetic beads were conjugated with either a 20 base poly-T oligo as illustrated in *i,* or with that oligo plus a 30 base poly-T oligo with an internal 5’-CAA-3’ sequence as illustrated in *ii*. Beads were extended using Duplase-3 and Cy5-ddGTP (* indicates unextended control during this step), after which, all samples were labeled with TdT and fluorescein-12-ddUTP. Oligos appear blue if extended with Duplase-3 and green if extended wtih TdT.

The ability of the Duplase to extend the single stranded oligonucleotides supports a dynamic transient hybridization model as illustrated in Fig. 1b, and intermolecular interactions alone are sufficient to facilitate extension of surface-bound oligos (Fig. 1c). To support this assertion, we have shown that oligo-dT sequences are not efficiently extended by the Duplase, because the oligo-dT is unable to hybridize intramolecularly. However, robust extension of an oligo-dT sequence is achieved when a second templating oligo is bound to the same surface (Fig. 1C).

### Polymerase-catalyzed cyclic reversible termination

Some DNA polymerases are able to catalyze the incorporation of a reversible terminator on a solid phase oligonucleotide. The templating strand may be the nascent strand itself through hairpin formation and cis-extension, a neighboring strand, or another oligo provided in solution (Fig. 2). Oligos cannot be extended beyond a single base addition due to the 3’ blocking group (3’-OR) on the newly added nucleotide. After the single base incorporation event, the 3’-hydroxyl is restored through a cleavage reaction. This process may be repeated for the addition of a second, third, etc. base. Chemically-cleavable reversible terminators, 3’-*O*-azidomethyl-dNTPs, were used, and deprotection was achieved using Tris (2-carboxyethyl) phosphine hydrochloride (TCEP), as first described by Palla *et al.* (20). However, we suspect that other reversible terminators could be used, such as chemically cleavable 3’-aminoxy groups (21), enzymatically cleavable(22), or UV-cleavable(23,24) reversible terminator nucleotides.

**Fig. 2.**
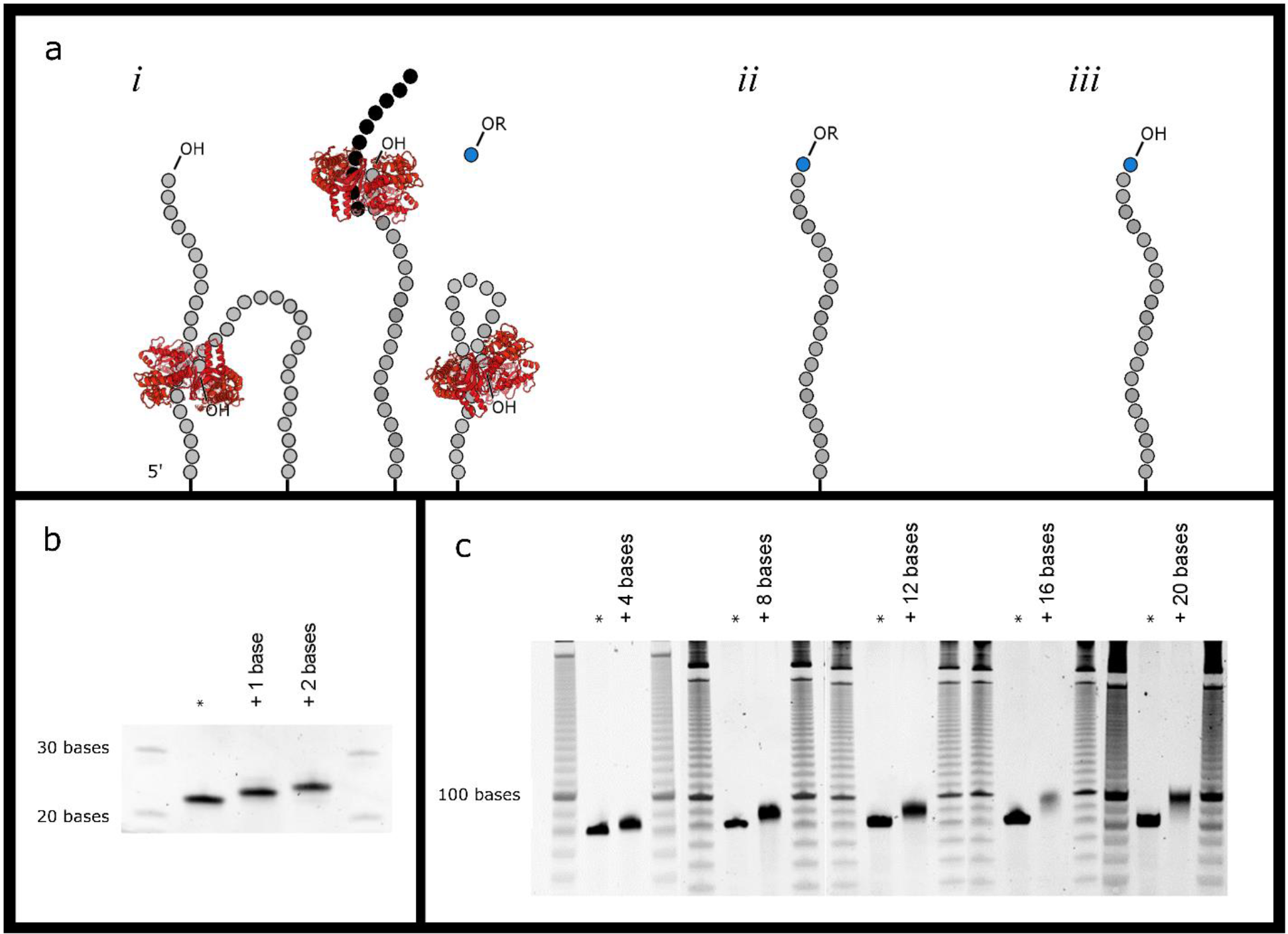
**(a) Polymerase-mediated solid-phase oligonucleotide synthesis using chemically blocked substrates.** DNA/RNA polymerases and RTs require a template for extension. (i) The templating strand for a surface-bound oligonucleotide may be a neighboring strand, an oligo in solution, or the nascent strand itself. A 3’-reversibly blocked nucleotide is added by the enzyme in solution to produce the oligo depicted in (ii), one base longer, but not capable of being extended. The 3’-OR group is then converted into a 3’-OH group through a cleavage step producing the oligo in (iii), which can be extended again. **(b) Sequential incorporation of two bases.** Denaturing PAGE analysis of a 20 base template, SPO-1, (after incorporation of 3’-O-azidomethyl-dTTP (+1 base), followed by cleavage of the 3’-O-azidomethyl capping group and incorporation of 3’-O-azidomethyl-dCTP (+ 2 bases). * indicates unextended control. **(c) Synthesis of a 20 base single-stranded DNA fragment on a universal templating oligo.** A 20 base sequence, ESO-1, was synthesized using this method on a universal templating oligo (UTO-1) using Duplase-3. Denaturing PAGE was performed after the addition of 4, 8, 12, 16, and 20 bases.

### The enzymes

Extension reactions using reversible terminators require enzymes with catalytic domains that are sufficiently large to accommodate the 3’-capping group. Many such enzymes have been reported, and numerous modified DNA polymerases have been developed for this purpose(25,26). Of the enzymes tested, the Moloney murine leukemia virus (M-MLV) RT, Superscript III, and the 9°N DNA polymerase, Therminator, are capable of extending solid-phase single stranded DNA with as few as two bases available for transient hybridization (Fig. 1a). Because the 9°N DNA polymerase has historically been the polymerase of choice for sequencing-by-synthesis experiments using 3’-O-azidomethyl reversible terminators(27,28), we generated a series of modified 9°N DNA polymerases variants to develop an optimized enzyme for EOS. We refer to our enzyme as a Duplase (available from Centrillion Biosciences).

Comparative analysis of the fidelity of three Duplase enzymes is shown in Fig. 3. Traditional primer extension reactions show that Duplase-1 is higher fidelity than Duplase-2, which is higher fidelity than Duplase-3 (Fig. 3a). Decreasing enzyme fidelity increases the probability of misincorporation events, thus increasing the chance that a base may be added to a surface-bound oligo when the oligo may exist in a conformation that does not necessarily template the base in solution since only one reversible terminator base is provided during each step of EOS (Fig.3b). This concept is illustrated in Fig. 3c, where a single oligonucleotide may rapidly switch between different transient hybridization states templating different bases. A high-fidelity polymerase will only incorporate A in one of the four structures drawn, whereas a low-fidelity polymerase might incorporate A opposite C or G or A as well. Therefore, the more promiscuous the enzyme, the higher the yield of the reaction product in the same amount of time. However, extension with a non-templated base (Fig. 3d) requires more time than extension with a templated base. In one minute, extension appears to be complete using the templated base, G, but no extension is observed using the non-templated base, A. If the reaction is allowed to proceed for one hour, however, extension can be observed using A. Since Duplase-3 has the lowest fidelity, we reason it will have the highest yield for EOS, and therefore it is used for the synthesis of the 20 base single-stranded oligonucleotide. Furthermore, Duplases are able to use ribonucleotides as a substrate (Supp. Fig. 1), and the 9°N polymerase has been used for XNA synthesis(29,30), increasing the options for substrates that can be synthesized using this technique.

**Fig. 3.**
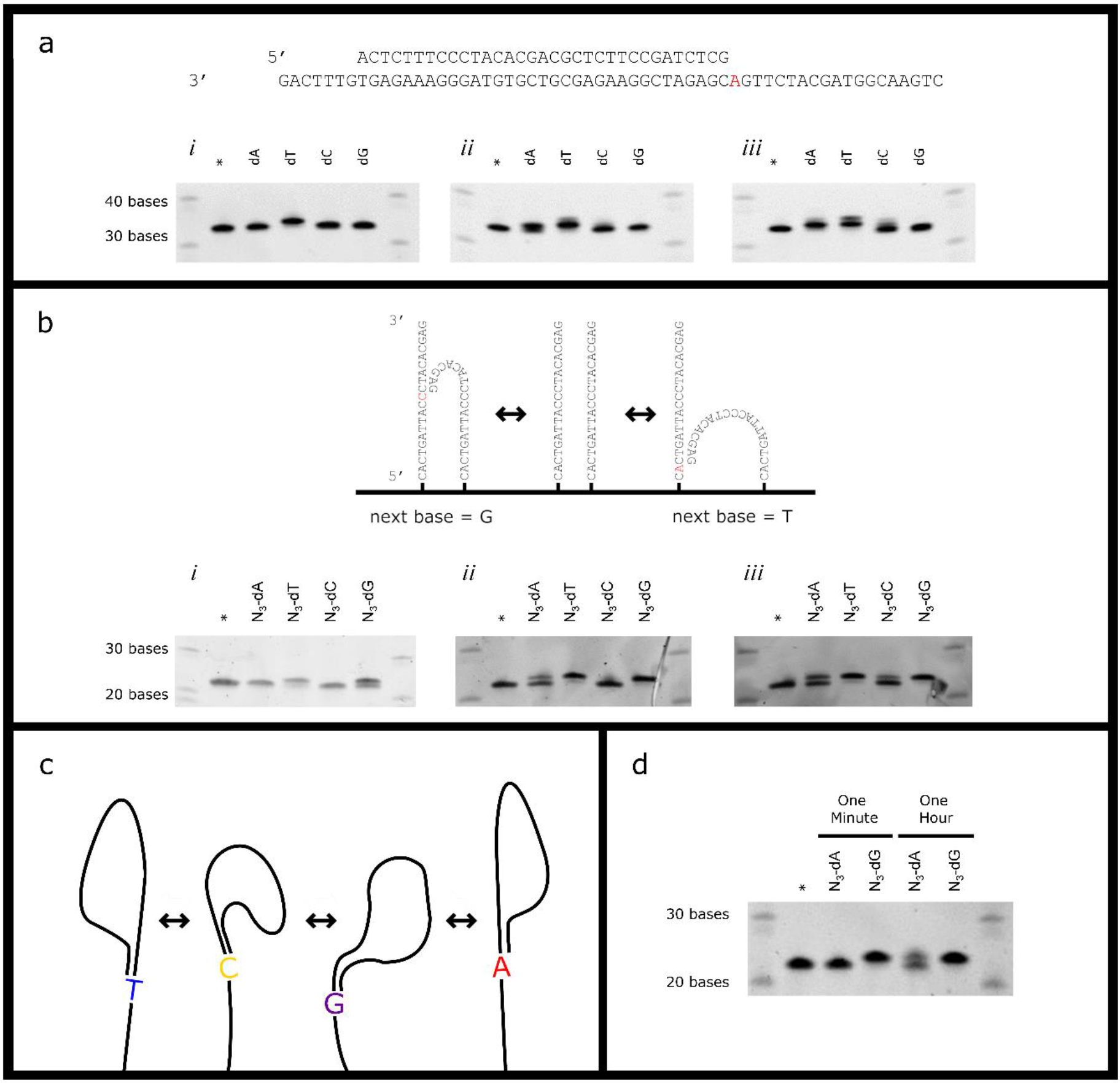
Effects of misincorporation on step-wise yield. **(a) Duplase fidelity on a double-stranded substrate.** Primer extension reactions were conducted using the two oligos sequences shown with the longer templating strand bound to streptavidin-coated magnetic beads through a 5’ biotin modification. The primer was stripped from the surface-bound template and analyzed by denaturing PAGE. Extension opposite adenine is shown with dATP, dTTP, dCTP, or dGTP used as the substrate in the reaction for (i) Duplase-1, (ii) Duplase-2, and (iii) Duplase-3. Increasing misincorporation can be seen. * indicates unextended primer control. **(b) Extension of a single-stranded substrate.** Reactions were conducted using a surface-bound single-stranded oligo, SPO-1, as shown with one of four 3’-O-azidomethyl-dNTP in solution and (i) Duplase-1, (ii) Duplase-2, and (iii) Duplase-3. * indicates unextended primer control. As polymerase fidelity decreases, the capability to incorporate bases not templated by a neighboring strand or hairpin structure increases. **(c) Illustration of transient hybridization.** An example oligonucleotide is shown templating four different bases for four different hybridization positions. Such an oligonucleotide at any point in time may exist in one of these conformations but not all of these conformations. **(d) Misincorporation reactions may be slower than templated reactions.**Extension of the single-stranded substrate in **(b)**, SPO-1, is demonstrated in one minute or one hour with Duplase-3 and 3’-O-azidomethyl-dATP (not templated by this oligonucleotide) or 3’-O-azidomethyl-dGTP (templated). In one minute, extension appears to be complete using the templated base, G, but no extension is observed using the non-templated base, A. If the reaction is allowed to proceed for one hour, however, extension can be observed using A.

### Optimization of reaction conditions

For EOS application, we want the enzyme to incorporate the provided base, regardless transient hybridization structures. Therefore, we assayed variables to increase the chance of any incorporation (which includes misincorporation) and found that reaction temperature and the divalent metal ion used had the most significant impact. Manganese has been shown to alter the geometry of the polymerase’s substrate binding pocket, opening it up to allow for polymerization with a non-templated base(31), and polymerases are temperature sensitive with respect to fidelity(32). Optimal extension of single-stranded oligos was achieved using manganese (Supp. Fig. 2) at 60°C (Supp. Fig. 3). Because the lifetime of hybridization structures decreases with increasing temperatures, these data support a model of transient DNA hybridization where oligonucleotides on a solid surface are rapidly flipping between hybridization states that may be recognized and extended by a DNA polymerase or RT.

### Extension opposite templates in solution

Random hexamer priming is a well-established method for cDNA synthesis, rolling circle amplification, and multiple displacement amplification for whole genome amplification(33,34). We sought to enhance extension of a surface-bound oligo by the addition of in-solution 3’-blocked oligonucleotides to serve as intermolecular hybridization partners for templated polymerase extension. It was determined that randomers increase incorporation in reactions with Duplase-3 and a short solid-phase primer (Fig. 4a). As with single-stranded oligos, reactions with random templates in solution are most efficient at higher temperatures, plateauing around 60°C (Supp. Fig. 3). Reaction yields are also higher with shorter randomers in solution (Supp. Fig. 4).

**Fig. 4.**
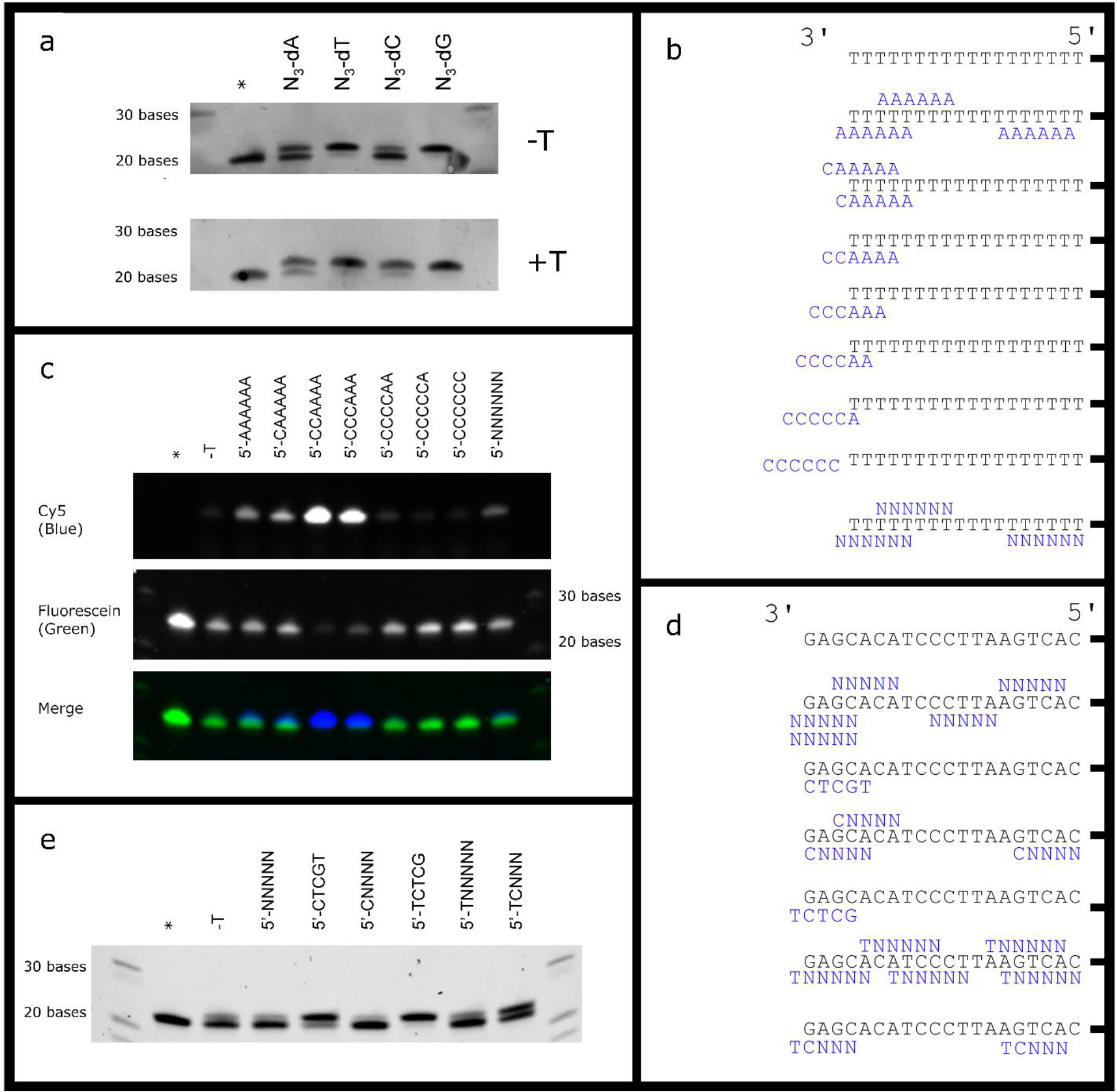
**(a) Extension of a short solid-phase oligo is enhanced by the addition of randomers to the reaction solution.** Denaturing PAGE analysis of a 20 base oligo extended with Duplase-3 and 3’-O-azidomethyl-dNTPs without added template to solution (−T) or with the addition of 3’-phosphate blocked random hexamers to the reaction solution (+T). * indicates unreacted control. **(b) Possible hybridization of different templates to a solid-phase poly-T sequence.** Templates used in (c) are drawn in blue. **(c) Three bases of hybridization are required with a template in solution.** Denaturing PAGE analysis of a poly-T sequence extended with Duplase-3 and Cy5-ddGTP. 3’-phosphate blocked hexamers are used in reaction solutions with the sequences listed. * indicates unreacted control. −T indicates reactions without added template. After extension with Duplase-3, all samples were incubated with TdT and fluorescein-12-ddUTP to label any primer that was not extended with Duplase-3. **(d) Possible hybridization of different templates to a solid-phase oligo.** Templates used in (e) are drawn in blue.Some potential sequence alignments during transient hybridization reactions are illustrated. **(e) Extension of a solid-216 phase oligo with specific, random, and semi-random templates in solution.** Denaturing PAGE analysis of SPO-1 after extension with Duplase-3 and 3’-O-azidomethyl-dATP. 3’-phosphate blocked pentamers are used in reaction 218 solutions with the sequences listed. * indicates unreacted control. −T indicates reactions without added template.

When the templating strand is in solution, Duplases require three bases for hybridization and at least one base overhang for efficient extension of a solid-phase primer (Fig. 4b-c). Increased base pairing requirements may result from instability that is not present when the template is a neighboring oligonucleotide tethered to the same solid surface. Non-random templates of known sequence are predictably more effective in increasing reaction yields that random templates are (Fig. 4d-e). There are, however, 1024 possible different sequences for a pentamer, the shortest oligonucleotide commercially available (4^5^). In order to develop a protocol for the synthesis of any possible sequence, all 1024 different pentamer sequences would have to be stocked and stored in separate wells in an automatic synthesizer, which is not feasible. Only one solution would be required for a random pentamer, NNNNN (N denotes a random base, A, T, C, or G). Four solutions would be required for a semi-random sequence, XNNNN, where X templates the incoming base, and 16 solutions would be required for the semi-random sequence, XYNNN, where Y is complementary to the last base of the solid-phase oligo. These two semi-random oligos improve synthesis yields over random pentamers when added to reaction solutions (Fig. 4d-e). However, there are obvious drawbacks to this method, for example when X=Y or for repetitive or homopolymeric stretches, and removal of templates from solution may be incomplete without additional purification steps and the use of templating oligos adds to the cost of the reactions. Instead, a universal templating oligo was used for the EOS of 20 bases as described below.

### Design of a universal template

A universal templating oligonucleotide (UTO) is an oligonucleotide sequence that can template any base regardless of the 3’ sequence. UTOs were originally designed including the universal base, 5-nitro-1-indolyl-3’-deoxyribose (5-NI)(35–37), however, there is no universal base that meets all desired requirements for EOS, ability to pair with all natural bases equally, prime DNA synthesis by DNA polymerases, and direct incorporation off each of the natural nucleotides(35,38). Therefore, we designed UTO-1, a 78 base sequence (Supp. Table 1), that contains all codons and includes poly-T sequences at the 3’ and 5’ ends that serve as spacers and a 3’ deoxyuracil. The 3’ deoxyuracil can be enzymatically cleaved using a combination of Uracil-DNA Glycosylase and apurinic/apyrimidic endonuclease to isolate the enzymatically synthesized fragment(39). All ANN, TNN, CNN, and GNN sequences are contained within UTO-1, meaning that any base can be templated during transient hybridization for efficient EOS. Using this approached, we synthesized the 20 base sequence ESO-1 (enzymatically synthesized oligo-1) on UTO-1 using Duplase-3. The resulting oligo was poly-adenylated, and adapters were added by PCR (Supp. Fig. 5), and the library was sequenced on a MiSeq sequencer (Illumina, San Diego, CA).

### Accuracy of the synthesized oligonucleotide

Background8 The sequencing data on the MiSeq was analyzed and 83,631 sequences of our synthesized oligonucleotide were obtained. These sequences were randomly divided into 10 groups for a statistical bootstrapped analysis of the incorporation efficiency and multiple base incorporation frequency (Fig. 5). The number of reads containing the full length ESO-1 sequence was compared to the number of reads one base shorter to determine the efficiency for the perfect incorporation of the final base, and these calculations were repeated to determine synthesis efficiency at each enzymatic synthesis step. Calculations excluded regions containing a homopolymer run because when the n−1 base was the same as the n base or the n+1 base, and therefore failure to synthesize either base would result in the same partial sequence. We observed that XX% of the sequences erroneously contained repeated bases. For example, some of the sequences containing N_13_CAG read as N_13_CAGG or N_13_CAGGG. Incorporation of the final base, C, in N_13_CAGC was thus calculated from the number of reads containing N_13_CAG less the number of reads containing N_13_CAGG. These incorrect sequences could be caused by either 3’OH contaminating bases in the extension mix (17) or TCEP carry over during the synthesis.

**Fig. 5.**
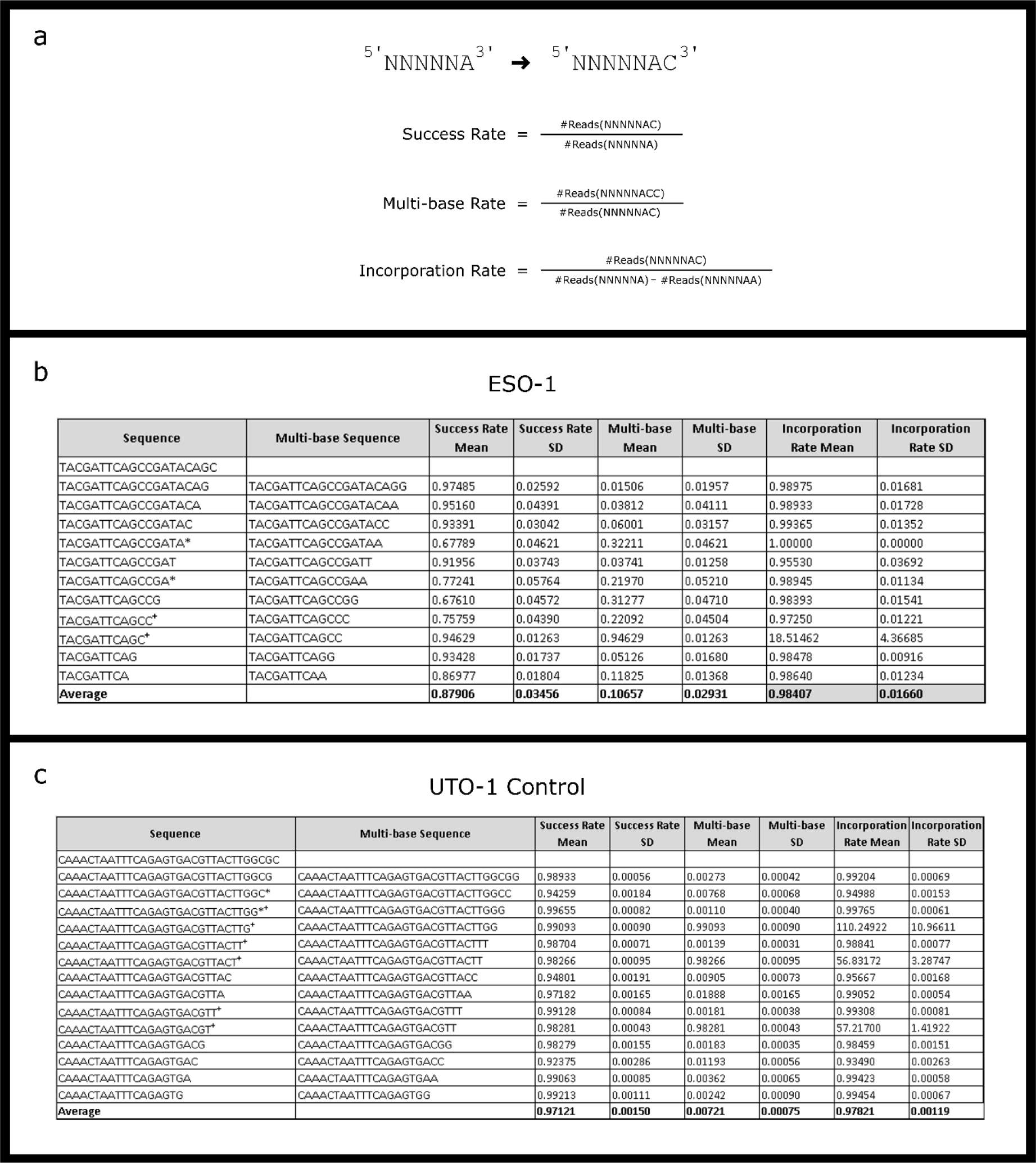
Oligo Synthesis Efficiency. **(a) Variables for evaluating efficiency.** The success rate, multiple base incorporation rate, and adjusted incorporation rate are calculated for the extension of the oligo NNNNNA with D. These numbers are calculated for the bootstrapped sequences for ESO-1 **(b)**, and for the partial UTO-1 sequence used as a control **(c)**. * indicates oligos for which the n+2 base is the same so that the sequence NNNNNAA as in the example in part (a) may be due to multi-base extension or C not being incorporated. + indicates oligos for which the n+1 base is the same so that multi-base extension cannot be distinguished from accurate synthesis. These numbers were omitted from analysis.

Based on the sequencing data, we calculated the stepwise efficiency for the last 12 bases of ESO-1 from the subset of sequences that correctly contained the first 8 bases. While the mean success efficiency for ESO-1 was 87.9%, after correcting for multiple base additions, the mean success efficiency was 98.4%. For the control sequence, the mean success efficiency was 97.1%, and incorporation efficiency was 97.8% (Fig. 5).

A secondary method of analyzing the stepwise incorporation efficiency was also implemented. The sequence alignment of the correct sequence and each MiSeq read was evaluated, as illustrated in Fig. 6 Each correctly aligned base was scored with a value of 1, and mismatches, insertions, and deletions were scored with 0. This allowed us to evaluate the accuracy of our final fragment, accounting for multiple addition events. The results indicated that 54.6% of ESO-1 reads achieved the maximum score of 19 (the first base of the 20 base sequence was omitted because it is the same as the last base on UTO-1), and 95.2% of the control sequence, UTO-1p2, reads achieved the maximum score of 14. This results in a step-wise efficiency of 98.6% and 99.6%, for ESO-1 and UTO-1p2 respectively.

**Fig. 6.**
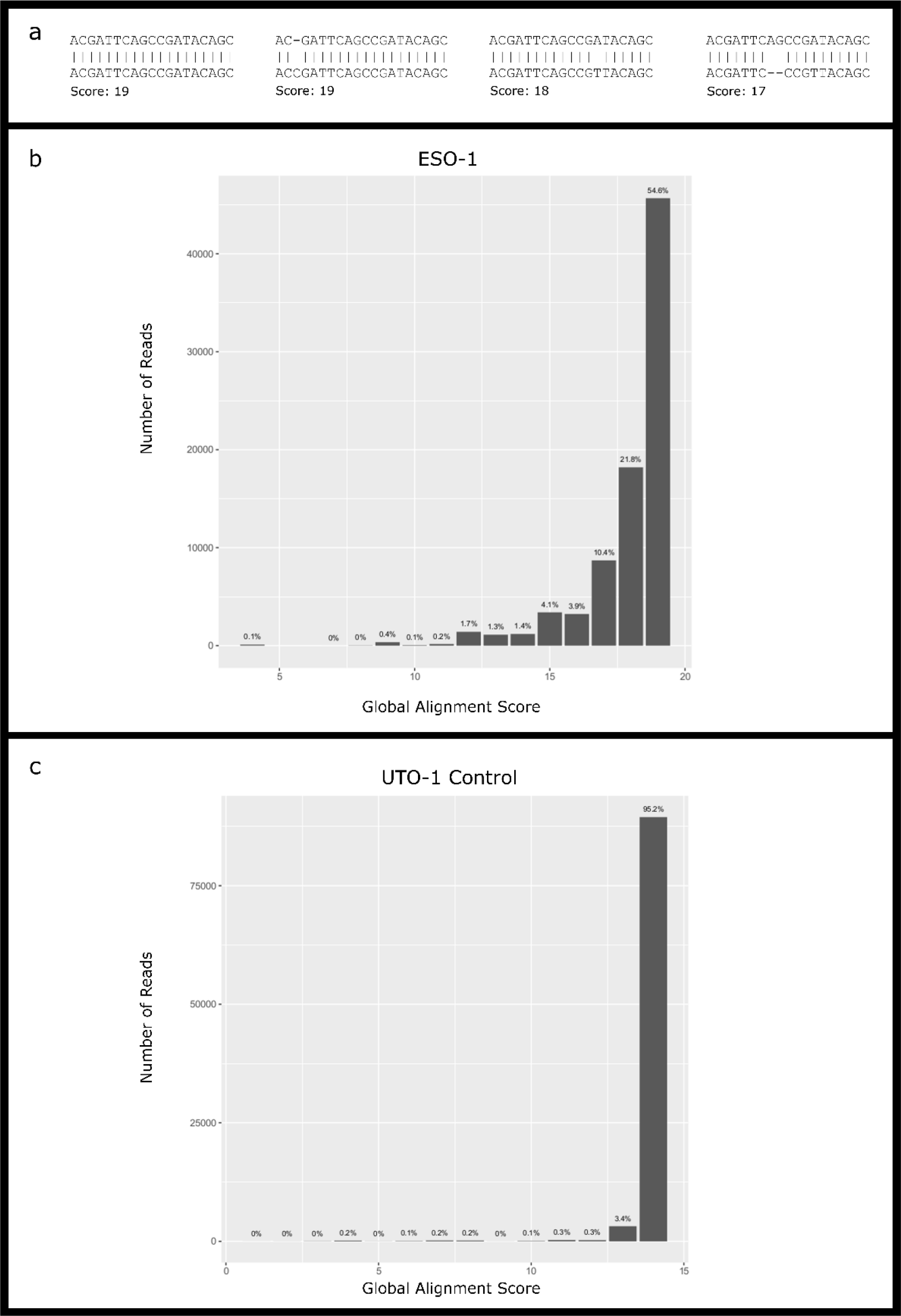
Read 1 Alignment Scores. **(a) Alignment Score Examples.** A value of 1 is assigned to every perfectly matched base. A maximum value of 19 was possible for ESO-1 (one base was omitted from analysis because the preceding sequence ended in the same base). A maximum value of 14 was possible for the PCR-prepared UTO-1p2 control sequence. Distribution of scores reflects final length of the correctly synthesized oligos accounting for multiple base incorporation, as displayed in **(b)** for ESO-1 and **(c)** for the UTO-1 control. 55.4% of sequences had a maximum value of 19 for ESO-1, and 95.2% of sequences had the maximum value of 14 for the UTO-1 control.

## Discussion

Since the isolation of the first DNA polymerase, *E. coli* pol I, by Kornburg in 1956, hundreds of other polymerases have been discovered, characterized, and engineered, and these polymerases have revolutionized the biomedical sciences with applications such as PCR and DNA sequencing. Herein, we have extended the capabilities of these enzymes and harnessed the same molecular machinery that made sequencing-by-synthesis possible to synthesize novel single-stranded DNA sequences. Enzymatic methods of DNA sequencing quickly overtook the Maxam-Gilbert chemical method more than 40 years ago; they were faster, less expensive, and able to produce longer reads. We hypothesize that DNA synthesis methods will follow a similar path with enzymatic methods opening the door for the production of longer, more accurate sequences at lower cost and without the production of hazardous waste.

We synthesized a 20 base single-stranded DNA oligo with stepwise yields between 98-99% using a DNA polymerase, reversible terminators, and a universal templating oligonucleotide. Our methods are already competitive with chemical methods, though further optimization will be required for the efficient synthesis of long oligos. For example, decreased enzymatic fidelity and mismatch discrimination would decrease reaction times and the length of the UTO required, and highly pure reversible terminators and a neutralization step following cleavage could eliminate multiple incorporation events. Additionally, we envision that the yield of full-length products without single-base deletions could be further increased by adding a capping step to terminate synthesis of unextended oligos after every step, and full-length products can be selected for by adding a final base either modified with a group that can be easily pulled-down such as biotin, or with one that may protect against exonucleolytic degradation in a cleanup step. It is reasonable to suspect that EOS can be improved to produce results greater than what is currently seen with chemical methods. High efficiency and low cost should encourage the use of this technology, leading to further advances.

## Methods

### Primer extension assays on double-stranded DNA

Primers were hybridized to a biotinylated target sequence immobilized on streptavidin-coated magnetic beads (Dynabeads MyOne Streptavidin T1, Thermo Fisher, Waltham, MA) by heating to 70°C for five minutes then 55°C for 15 minutes then 25°C for five minutes in RB (1M sodium chloride, 25mM Tris-HCl pH 7.5, 0.01% TWEEN20). 0.1mg of beads were used per 25μL reaction. Reactions were started by the addition of nucleotide at temperature and stopped by the addition of a 1.6x volume of RB. The hybridized primer was stripped from the bead-bound template using 0.1N sodium hydroxide for ten minutes at room temperature and analyzed by denaturing PAGE. All oligos were purchased from Integrated DNA Technologies (IDT, Redwood City, CA). Sequences are listed in Supp. Table 1.

### Preparation of single-stranded DNA on a solid surface for extension

Single-stranded oligos were immobilized to streptavidin coated beads. 0.1mg of beads were used per 25μL reaction volume. Reactions were started by the addition of nucleotide at temperature and stopped by the addition of a 1.6x volume of RB. The immobilized primer was stripped from the bead-bound template in 0.1N sodium hydroxide for five minutes at 65°C and analyzed by denaturing PAGE. Oligos were purchased from IDT and are listed in Supp. Table 1.

### Primer extension reactions with Duplase enzymes

For reactions on double-stranded DNA, magnetic beads were prepared as previously described with immobilized template and hybridized primer. Beads were washed in reaction buffer and then resuspended in the reaction mix containing a final concentration of 20mM Tris-HCl pH8.8, 10mM ammonium sulfate, 10mM KCl, 0.1% Triton X-100, 2mM MgSO_4_, 1mg/mL bovine serum albumin (Sigma-Aldrich, St. Louis, MO), 4μg/mL polyvinylpyrrolidone 10, and 1ug of enzyme. Reaction mixtures were pre-warmed to 45°C, and nucleotide was added to a final concentration of 2μM. Reactions were allowed to proceed for one minute before stopping for analysis by denaturing PAGE. Extension reactions on single-stranded DNA were performed as previously described though buffer components, reaction times and temperatures, and nucleotide concentrations varied across experiments conducted for research purposes. Optimized conditions for the synthesis of the 20 base ESO-1 sequence are described below.

### Extension reactions with commercially available enzymes

Extension reactions on single-stranded DNA were performed similarly to the reactions with Duplase enzymes described above for 60 minutes at 60°C using 2U Vent (NEB, Ipswich, MA), 2U Deep Vent (NEB), or 2U Therminator (NEB) in 20mM Tris-HCl pH8.8, 10mM ammonium sulfate, 10mM KCl, 0.1% Triton X-100, 8mM manganese chloride, 10uM dGTP. Extension reactions with 2U Phusion (NEB) were performed for 60 minutes at 60°C using Phusion high fidelity buffer, 8mM manganese chloride, 10uM dGTP. Extension reactions with 10U Phi29 (NEB) were performed for 60 minutes at 37°C using Phi29 buffer, 8mM manganese chloride, 10uM dGTP. Extension reactions with 5U Klenow (NEB) or 3U T4 DNA polymerase (NEB) were performed for 60 minutes at 37°C using NEBuffer 2, 8mM manganese chloride, 10uM dGTP. Extension reactions with 250U Superscript III (Thermo Fisher) were performed for 5 minutes at 25°C, 5 minutes at 50°C, then 30 minutes at 55°C using first strand buffer, 8mM manganese chloride, 10uM dGTP. Reactions were analyzed by denaturing PAGE.

### Cleavage of Reversible Terminators

After incorporation of reversible terminators, beads were resuspended in 50mM TCEP pH9.0 (Gold Bio, Olivette, MO) and incubated at 60°C for 10 minutes. The TCEP solution was removed, and beads were resuspended in RB and transferred to a new reaction vessel.

### Synthesis of the 20 base ESO-1 sequence with SPO-13

Beads were prepared as previously described with immobilized SPO-13. 0.2mg of beads were used in 50μL reactions containing 200mM NaCl, 20mM Tris pH 8.0, 8mM MnCl_2_, 1μg Duplase-3, and 100μM of a 3’-*O*-azidomethyl-dNTP. Reactions were allowed to proceed for one hour before stopping. The sequence ESO-1 was synthesized by the incorporation of 3’-*O*-azidomethyl-dCTP followed by cleavage with TCEP and the incorporation of 3’-*O*-azidomethyl-dGTP, followed by cleavage with TCEP, etc. For the first synthesis, after the addition of 4, 8, 12, 16, and 20 bases, 0.025mg of beads was removed from solution, and the oligo was stripped from beads and analyzed by denaturing PAGE. Volumes for next steps were adjusted accordingly. After the final incorporation/cleavage step, the oligo with ESO-1sequence was used for sequencing.

### Fluorescent labeling of oligos with terminal deoxynucleotidyl transferase (TdT)

Labeling of poly-T sequences, which do not stain well with SYBR Gold, was accomplished by end-labeling oligos with TdT (New England Biolabs, Ipswich, MA) and fluorescein-12-ddUTP (Perkin Elmer, San Jose, CA) by incubating 0.1mg of beads with 5μM nucleotide and 20U of enzyme in 20mM Tris-HCl pH7.5, 10mM ammonium sulfate, 10mM KCl, 0.1% Triton X-100, and 0.5mM MnCl_2_. Reactions were allowed to proceed for 3 hours at 37°C before heat-inactivation of the enzyme for 20 minutes at 75°C.

### Sequencing of SPO-13 with synthesized 20 base AM1 sequence

The SPO-13-AM1 oligo still on beads was poly adenylated using TdT (Roche, Santa Clara, CA) and Duplase-1. Illumina sequencing adapters were added to these sequences using PCR, a poly (T)-tailed P7 adapter sequence, P7-Poly (T), and an oligo with the sequence PCR-ESO1-FPCR was repeated with Illumina P5 and P7 adapter oligos, and the library quality was analyzed by Bioanalyzer (Agilent, Santa Clara) and qPCR. All oligos were purchased from IDT. A MiSeq nano flow cell (Illumina) was clustered with a 4pM library comprised of 90% PhiX and 10% PCR-prepared library. 150 step paired end cycling was run using a V2 kit (Illumina), and the AM1 sequence was identified by manually sorting first output reads. Background noise was filtered by selecting for part of the universal templating oligonucleotide sequence, UTO-1p (Supp. Table 1)

### Denaturing PAGE

Samples were resolved by electrophoresis in 6% or 15% polyacrylamide TBE-Urea gels (Thermo Fisher) in TBE. Gels were stained in SYBR Gold (Thermo Fisher) for five to ten minutes at room temperature and imaged on a ChemiDoc MP system (Bio-Rad, Hercules, CA). Fluorescent products were visualized before and after staining.

## Supporting information

Hoff EOS Supp

## Acknowledgements

We would like to thank Justin Costa and Janet Warrington for their help and expertise writing. We would also like to thank Foster Hoff for his help in designing UTO-1 and Jiayi Sun for her help in analyzing sequencing data.

