## Supplementary material for "Rapid and dynamic nucleic acid hybridization enables enzymatic oligonucleotide synthesis by cyclic reversible termination: A novel mechanism for enzymatic DNA synthesis": Hoff EOS Supp

| Oligonucleotide Name | Sequence 5'-3' | Use |
|---|---|---|
| UTO-1 | /5Biosg/TTTTTACCTCTGTATGCATCGATAGGTCCGGGAAGCCCACAAACTAATTTCAGAGTGACGTTACTTGGCGCTTTTTT/3deoxyU/ | Single-stranded DNA Extension |
| ESO-1 | TACGATTCAGCCGATACAGC | Synthesized with Duplase-3 |
| UTO-1p | CAAACTAATTTCAGAGTGACGTTACTTGGCGCTTTTTTT | Sequencing analysis |
| UTO-1p2 | ACGTTACTTGGCGC | Sequencing analysis |
| SPO-1 | /5Biosg/CACTGATTACCCTACACGAG | Single-stranded DNA Extension |
| SPO-2 | /5Biosg/GTGACTAATGGGATGTGCTC | Single-stranded DNA Extension |
| SPO-3 | /5Biosg/CACTGATTTCTCTACACGAG | Single-stranded DNA Extension |
| SPO-4 | /5Biosg/CACTGATTAGGCTACACGAG, | Single-stranded DNA Extension |
| SPO-30bp | /5Biosg/CTGAACGGTAGCATCTTGACGAGATCGGAAGAGCGTCGTGTAGGGAAAGAGTGTTTCAG | Single-stranded DNA Extension |
| T18-Biot | /5Biosg/TTTTTTTTTTTTTTTTTT | Single-stranded DNA Extension |
| T20-Biot | /5biosg/TTTTTTTTTTTTTTTTTTTT | Single-stranded DNA Extension |
| T7CAAT20-Biot | /5Biosg/TTTTTTTCAATTTTTTTTTTTTTTTTTTTT | Single-stranded DNA Extension |
| A6-P | AAAAAA/3Phos/ | In-solution Templates |
| CA5-P | CAAAAA/3Phos/ | In-solution Templates |
| C2A5-P | CCAAAA/3Phos/ | In-solution Templates |
| C3A3-P | CCCAAA/3Phos/ | In-solution Templates |
| C4A2-P | CCCCAA/3Phos/ | In-solution Templates |
| C5-A-P | CCCCCA/3Phos/ | In-solution Templates |
| C6-P | CCCCCC/3Phos/ | In-solution Templates |
| N6-P | NNNNNN/3Phos/ | In-solution Templates |
| N9-P | NNNNNNNNN/3Phos/ | In-solution Templates |
| N12-P | NNNNNNNNNNNN/3Phos/ | In-solution Templates |
| N15-P | NNNNNNNNNNNNNNN/3Phos/ | In-solution Templates |
| N18-P | NNNNNNNNNNNNNNNNNN/3Phos/ | In-solution Templates |
| N21-P | NNNNNNNNNNNNNNNNNNNNN/3Phos/ | In-solution Templates |
| N5-P | NNNNN/3Phos/ | In-solution Templates |
| CTCGT-P | CTCGT/3Phos/ | In-solution Templates |
| CN4-P | CNNNN/3Phos/ | In-solution Templates |
| TCTCG-P | TCTCG/3Phos/ | In-solution Templates |
| TN5-P | TNNNNN/3Phos/ | In-solution Templates |
| TCN3-P | TCNNN/3Phos/ | In-solution Templates |
| ds-template | 5Biosg/CTGAACGGTAGCATCTTGACGAGATCGGAAGAGCGTCGTGTAGGGAAAGAGTGTTTCAG | Primer Extension |
| ds-primer1 | ACTCTTTCCCTACACGACGCTCTTCCGATCTCG | Primer Extension |
| ds-primer2 | CACTCTTTCCCTACACGACGCTCTTCCGATCTC | Primer Extension |
| PCR-ESO1-F | TATGCATCTACACTCTTTCCCTACACGACGCTCTTCCGATCTTTTTTACCTCT*G. | Sequencing Library Preparation |
| P7-Poly(T) | CTCGGCATTCCTGCTGAACCGCTCTTCCGATCTTTTTTTTTTTTTTTTTTTTT*T | Sequencing Library Preparation |

**Supp. Table 1. Oligo Sequences.** All DNA sequences used for extension reactions, sequencing library preparation, and sequencing analysis are listed here. Modifications (5'-biotinylation, 3'-deoxyuracil, 3'-phosphates, and thio bonds) are listed according to ordering specifications for IDT.**Supp. Fig. 1. M-MLV Reverse Transcriptases (RTs) can incorporate dNTPs but not reversible terminators on single-stranded DNA. (a) DNA sequences tested for extension with M-MLV RTs.** Extension was assayed using (i) a 20 base single-stranded solid phase oligo, or (ii) a 33 base primer hybridized to a template. Both sequences template incorporation of guanosine. **(b) Extension with Superscript IV.** (i) Superscript IV can incorporate dGTP but not 3'-*O*-azidomethyl-dGTP on single-stranded DNA. (ii) Using the same reaction conditions, extension can be seen with both nucleotides on double-stranded DNA. **(c) Extension with SMARTScribe.** (i) SMARTScribe can incorporate dGTP but not 3'-*O*-azidomethyl-dGTP on single-stranded DNA. (ii) Using the same reaction conditions, extension can be seen with both nucleotides on double-stranded DNA.

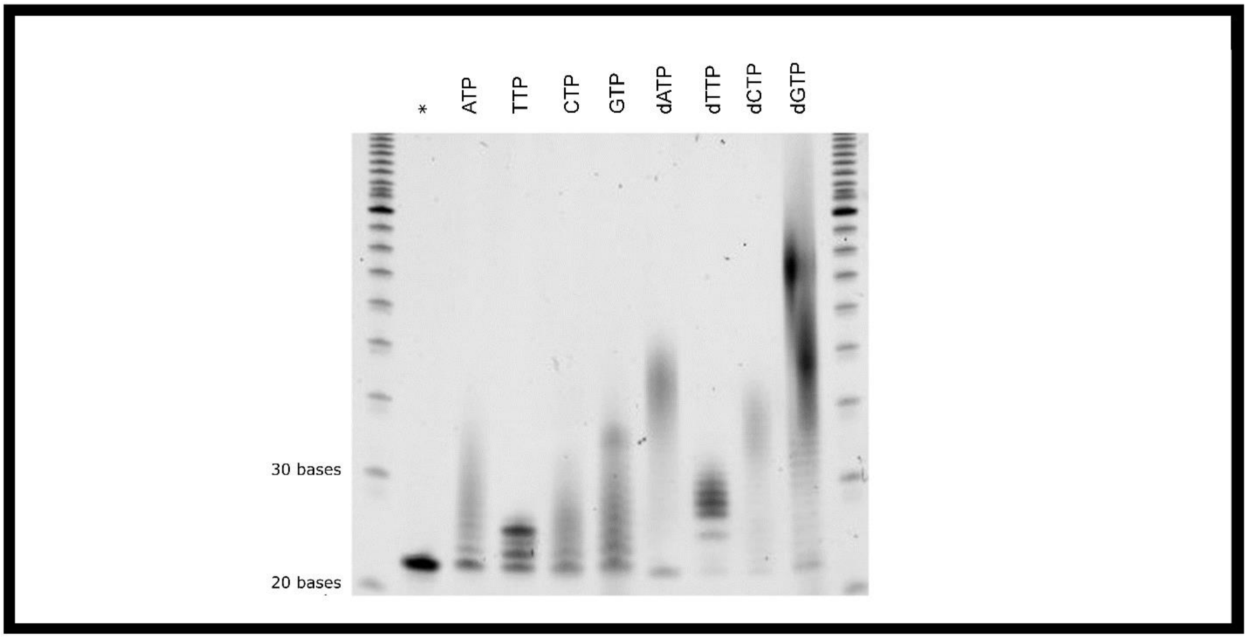


**Supp. Fig. 1. Duplase-3 can incorporate ribonucleotides on a single-stranded oligo.** Denaturing PAGE analysis of a 20 base single-stranded surface-bound oligo, SPO-1, extended with Duplase-3 and NTPs or dNTPs. * indicates unextended control.

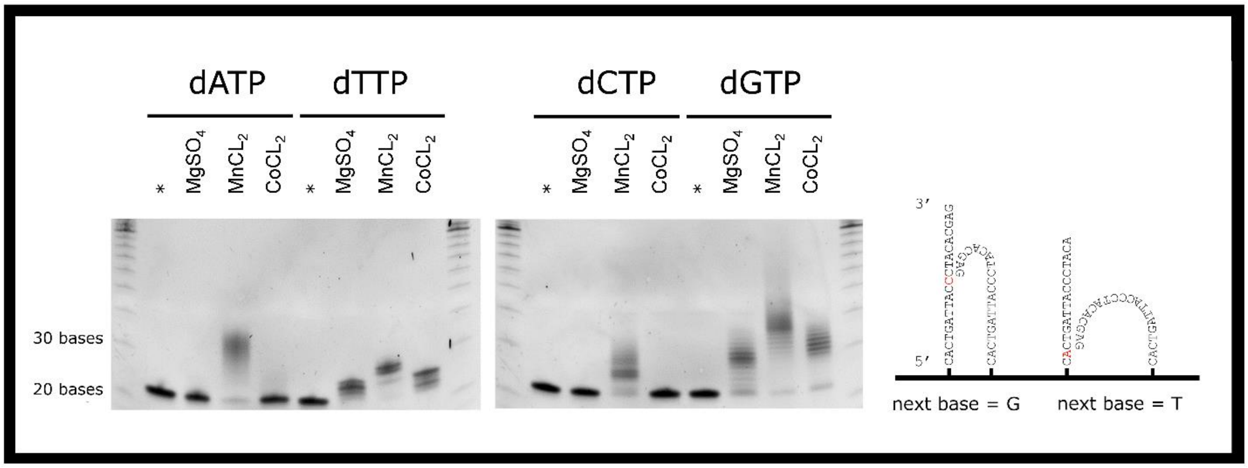


**Supp. Fig. 2. Effect of different divalent metals on single-stranded oligo synthesis.** Denaturing PAGE analysis of a 20 base single-stranded surface-bound oligo extended with Duplase-3 and dATP, dTTP, dCTP, and dGTP using magnesium, manganese, or cobalt as a cofactor in solution. Extension of templated bases is achieved with all three metals, but extension of non-templated bases is only achieved with manganese.

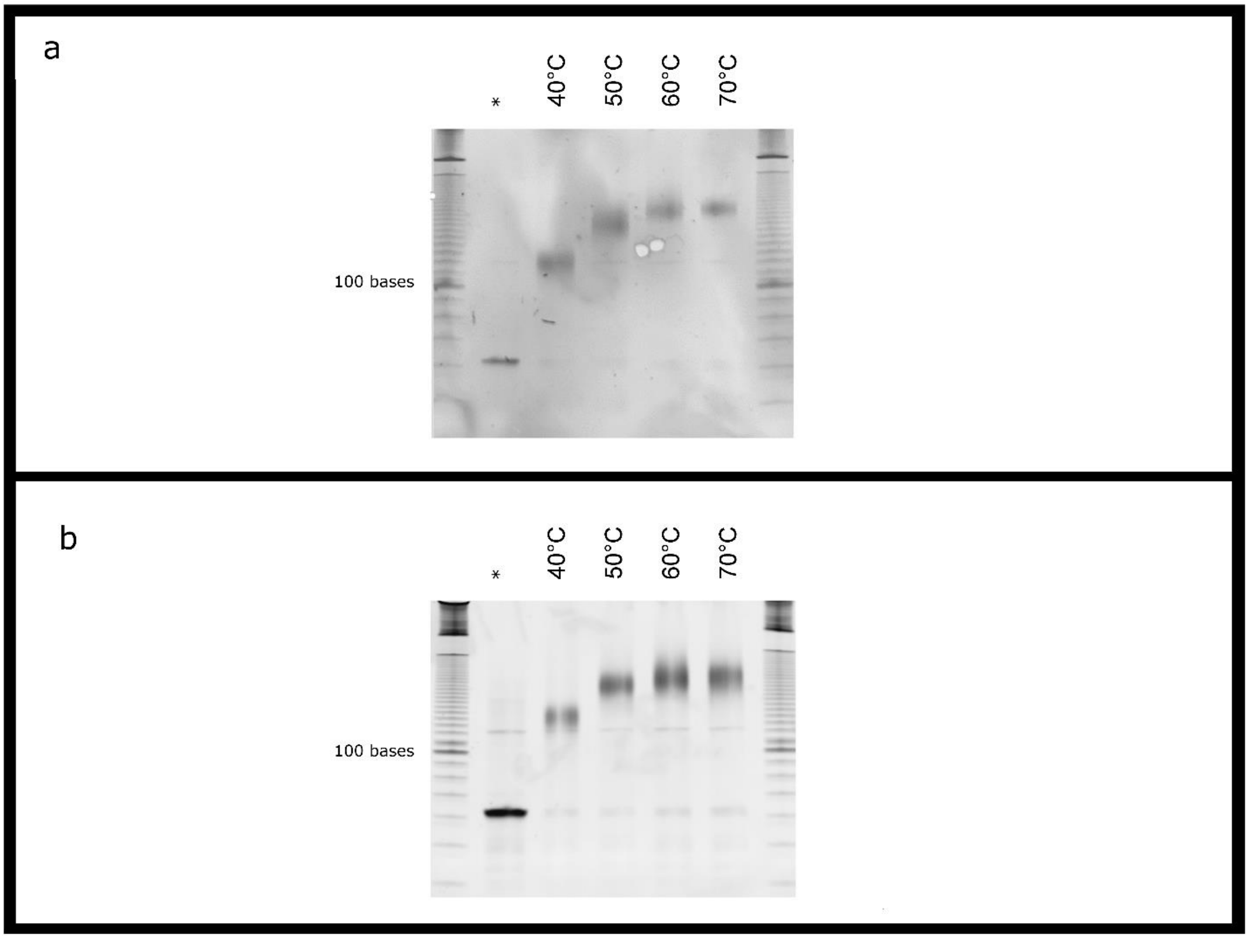


**Supp. Fig. 3. Temperature effects on single-stranded extension with Duplase-3.** Denaturing PAGE analysis of a 59 base solid-phase oligo extended with Duplase-3 and dCTP at different reaction temperatures. Reactions were executed with **(a)** the absence of a template in solution, or **(b)** 3'-phosphate blocked random hexamer templates in solution.

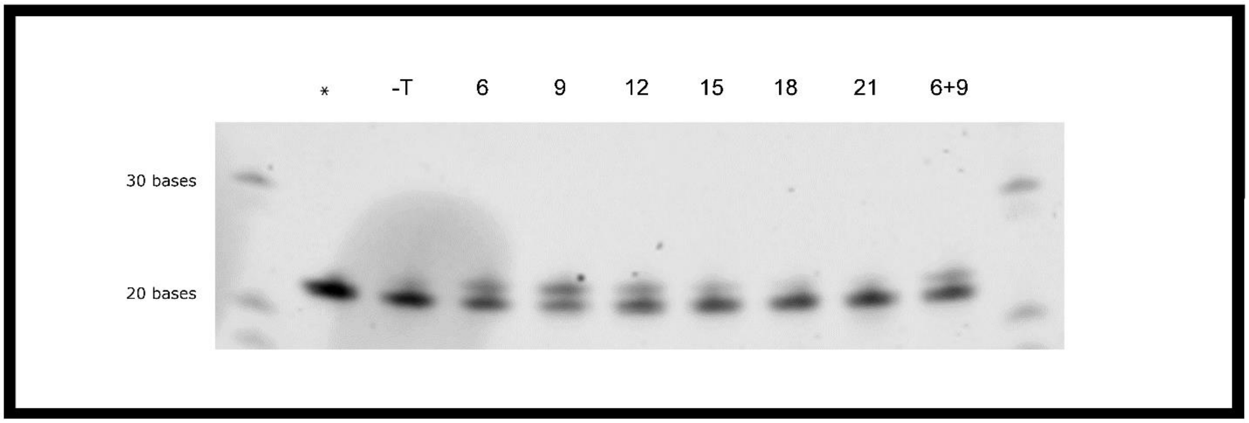


**Supp. Fig. 4. Effects of randomer length.** Denaturing PAGE analysis of a 20 base solid-phase oligo extended with Duplase-3 and 3'-*O*-azidomethyl-dATP with 3'-phosphate blocked random templates in solution. Randomers 6, 9, 12, 15, 18, 21, or 6 and 9 bases in length were added to the reaction mix. * indicates unextended control. -T indicates control sample without in-solution template added.

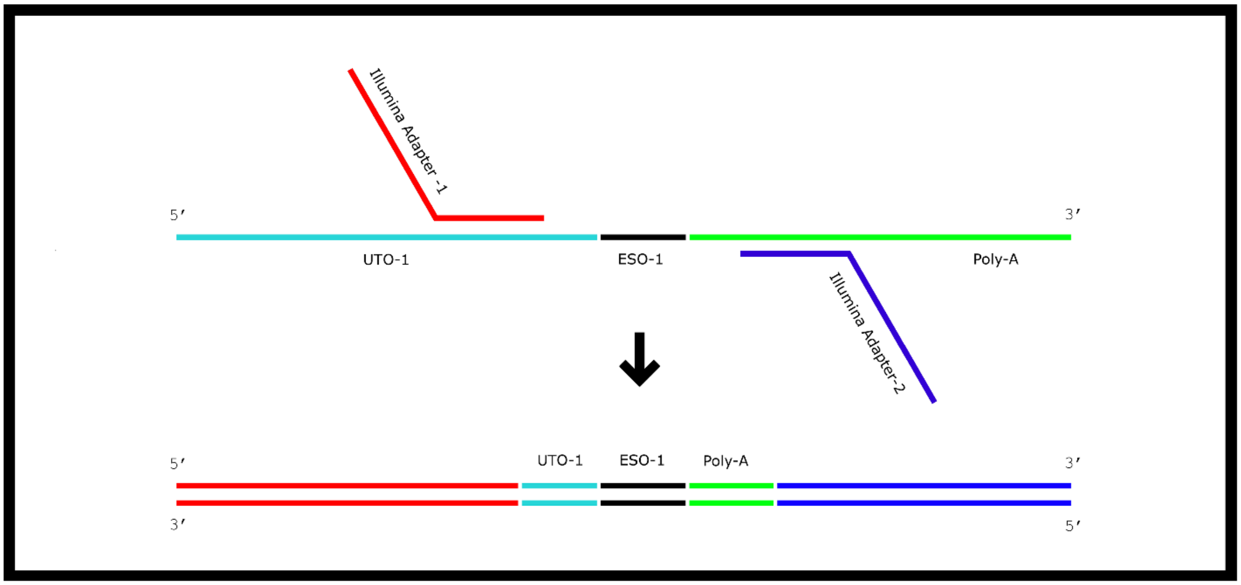


**Supp. Fig. 5. Library Prep of ESO-1.** An Illumina sequencing library was prepared from the ESO-1 sequence on UTS-1 by poly-adenylation of the oligo followed by two rounds of PCR. The first round of PCR added partial Illumina adapters as illustrated using tailed PCR primers with partial sequence identity to UTS-1 on the first primer and a poly-T tail on the second primer. The second round of PCR used non-indexed full-length Illumina primers. The resulting PCR product contained a partial UTS-1, ESO-1, and a poly-A tail sandwiched between adapters.
